# Room-temperature-storable PCR Mixes for SARS-CoV-2 Detection

**DOI:** 10.1101/2020.04.07.029934

**Authors:** Jiasu Xu, Jin Wang, Zecheng Zhong, Xiaosong Su, Kunyu Yang, Zhongfu Chen, Dongxu Zhang, Tingdong Li, Yingbin Wang, Shiyin Zhang, Shengxiang Ge, Jun Zhang, Ningshao Xia

**Affiliations:** State Key Laboratory of Molecular Vaccinology and Molecular Diagnostics, National Institute of Diagnostics and Vaccine Development in Infectious Diseases,School of Public Health, Xiamen University, Xiamen 361102, China; Xiamen International Travel Healthcare Center, Xiamen 361013, China

**Keywords:** COVID-19, SARS-CoV-2, freeze-drying, PCR

## Abstract

A novel coronavirus (severe acute respiratory syndrome coronavirus 2, SARS-CoV-2) emerged in late 2019, causing an outbreak of pneumonia [coronavirus disease 2019 (COVID-19)] in Wuhan, China, which then rapidly spread globally. Although the use of ready-made reaction mixes can enable more rapid PCR-based diagnosis of COVID-19, the need to transport and store these mixes at low temperatures presents challenges to already overburdened logistics networks. Here, we present an optimized freeze-drying procedure that allows SARS-CoV-2 PCR mixes to be transported and stored at ambient temperatures, without loss of activity. Additive-supplemented PCR mixes were freeze-dried. The residual moisture of the freeze-dried PCR mixes was measured by Karl-Fischer titration. We found that freeze-dried PCR mixes with ∼1.2% residual moisture are optimal for storage, transport, and reconstitution. The sensitivity, specificity, and repeatability of the freeze-dried reagents were similar to those of freshly prepared, wet reagents. The freeze-dried mixes retained activity at room temperature (18∼25°C) for 28 days, and for 14 and 10 days when stored at 37°C and 56°C, respectively. The uptake of this approach will ease logistical challenges faced by transport networks and make more cold storage space available at diagnosis and hospital laboratories. This method can also be applied to the generation of freeze-dried PCR mixes for the detection of other pathogens.

## 1 Introduction

The current coronavirus disease 2019 (COVID-19) outbreak caused by severe acute respiratory syndrome coronavirus 2 (SARS-CoV-2) is a public health emergency of international concern^[1, 2]^. At the time of writing (6th April 2020), at least 207 countries have been affected, with at least 1 210956 cases and 67 594 deaths globally^[3]^. Both infected persons and asymptomatic carriers of SARS-CoV-2 are likely sources of new infections^[4, 5]^. Timely diagnosis and management are essential for disease control. Real-time reverse transcriptase-polymerase chain reaction (rRT-PCR) is an accurate and sensitive molecular technique and is considered the “gold standard” for the diagnosis of COVID-19^[6, 7]^.

However, to maintain bioactivity, PCR reagents must be transported and stored at a low temperature. This presents challenges to already overburdened transport logistics networks and cold storage space at diagnosis and hospital laboratories.

Freeze-drying (lyophilization) is a low-temperature dehydration process mainly used for stabilizing of heat-labile biological drug substances contained in aqueous solutions^[8]^. Because water drives many destabilization pathways, removing most of the water can prolong the shelf-life of the product^[9, 10]^. Because freeze-dried reagents typically contain all of the necessary components for testing (at appropriate concentrations), errors associated with improper handling of wet reagents can also be reduced. This reduces preparation time and, thus, testing throughput.

There have been several recent publications investigating the possibility of freeze-drying PCR mixes. Klatser et al. freeze-dried PCR mixes for the detection of mycobacterium, which could be stored at 4°C and 20°C for 1 year and at 56°C for 1 week^[11]^. Tomlinson et al. freeze-dried PCR mixes for the detection of *Phytophthora ramorum*, which could be stored at room temperature for 20 weeks.^[12]^ Takekawa et al. freeze-dried a PCR mix for the detection of avian influenza virus in wild birds, but did not report the preservation time^[13, 14]^.

However, there are some important challenges associated with the freeze-drying of PCR mixes that have not yet been adequately addressed. Efforts to lyophilize PCR mixes for the detection of RNA virus are complicated by the instability of reverse transcriptase^[15, 16]^. Klatser et al.^[11]^ and Tomlinson et al.^[12]^ did not include a reverse transcriptase in their PCR mixes. Although Takekawa et al. targeted an RNA virus, they did not report long-term stability test or accelerated stability test data^[13, 14]^. This is particularly relevant to the current study given that SARS-CoV-2 is a single-stranded RNA coronavirus^[17]^.

Physical evaluation methods are critical when developing storable molecular biology tools, but the published works have often neglected this. For example, the residual moisture content is the most important factor affecting the quality and stability of freeze-dried reagents^[18, 19]^, and the commonly applied reduced weight method is inadequate. Karl-Fischer (KF) titration is an absolute method for measuring residual moisture content and is accepted as the standard method for water content determination in freeze-dried reagents^[20]^, but is rarely applied in studies because of its complexity.

In addition, the choice of assessment method to evaluate the freeze-dried reagents is pivotal. For example, regular PCRs are not quantitative, whereas rRT-PCR can report the dynamic changes in product abundance during the whole process, and can be used to detect reaction inhibitors or reduced activity.

Here, we propose a methodology for freeze-drying PCR mixes for the detection of SARS-CoV-2. Multiple physical assessment methods, such as Karl-Fischer titration and appearance evaluation, have been applied. To better assess the detection performance of the freeze-dried PCR reagents, we have used rRT-PCR to test samples gathered at the Xiamen International Travel Healthcare Center. We compare the sensitivity, specificity, and repeatability between the freeze-dried reagents and the wet reagents with consistent results. The freeze-dried reagents are thermostable and can be store at room temperature, 37°C, or 56°C for lengthy periods.

## 2 Materials and methods

### 2.1 Clinical specimens

Twenty-six clinical throat swab specimens were collected at the Xiamen international travel healthcare center. Five of these were from patients who had been diagnosed as having COVID-19. The collected specimens were stored in a 1.5-ml sample freezer tube and maintained at -80°C before nucleic acid extraction. RNA was extracted using the DOF-9648 purification system (GenMagBio, China) according to the manufacturer’s protocol.

### 2.2 rRT-PCR

The 40-μL reactions contained 5 μL of RNA, 0.4 μL of TAKARA Taq™ Hot Start Version (TAKARA, Japan), 4 μL of 10 × PCR Buffer (Mg^2+^ plus) provided with the TAKARA Taq™ Hot Start Version (TAKARA, Japan), 0.08 μL of *TransScript*® Reverse Transcriptase [M-MLV, RNaseH-](TransGen Biotech, China), 4 μl of 2.5 mM of each deoxyribose triphosphates (dNTPs) (TAKARA, Japan), and 1 μl of 10 mM of primers or TaqMan probes.

The primers and probes were designed according to the open reading frames of the genes encoding the 1ab (ORF1ab), nucleocapsid (N), and spikes (S) proteins of SARS-CoV-2. We downloaded these sequences from GenBank, and designed the related primers and probes using Mega version 7 and Oligo version 6 software. All oligonucleotides were synthesized and provided by Sangon Biotech (Shanghai, China) (Table 1).

**Table 1.**
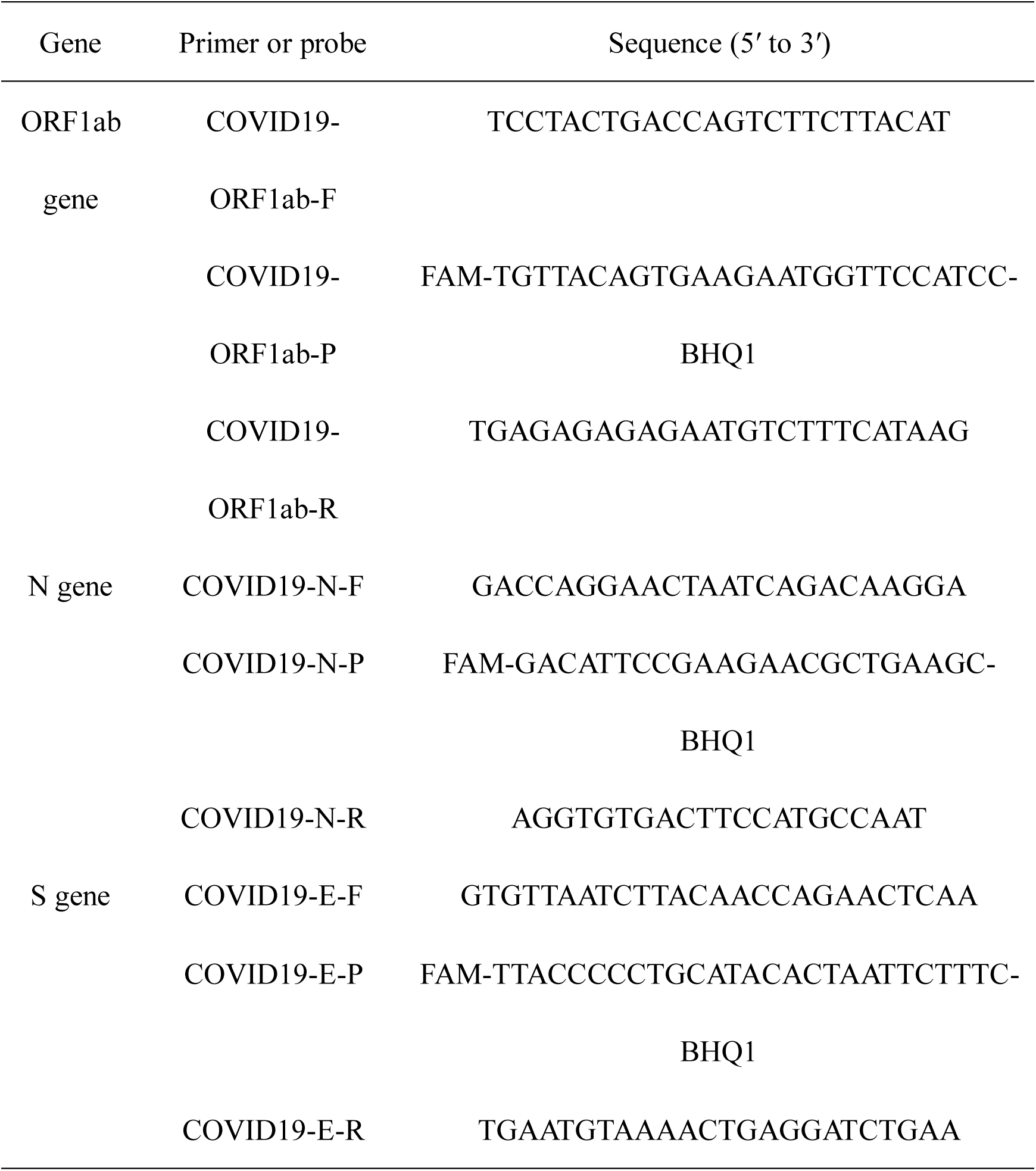
The primers and TaqMan probes used in this study.

Thermal cycling was performed at 50°C for 5 min for reverse transcription, followed by 95°C for 10 min and then 40 cycles of 95°C for 15 s, and 55°C for 30 s. All rRT-PCR assays were done using a CFX96 Touch instrument (CT022909, Bio-Rad, USA).

### 2.3 Freeze-drying

The PCR mixes were supplemented with trehalose [10% final concentration (w/v), Sigma-Aldrich], mannitol [1.25% final concentration (w/v), Sigma-Aldrich], BSA [0.002% final concentration (w/v), TAKARA] and polyethylene glycol 20000 (PEG20000) [0.075% final concentration (w/v), Sigma-Aldrich]. Then, the mixes were aliquoted into PCR tube strips (TCS-0803, Rio-Rad) before freeze-drying.

The freeze-drying process consists of multiple consecutive phases. First, we loaded the PCR tube strip containing the reagents into the shelf of the freeze dryer (Advantage 2.0,VITRIS), then lowered the shelf temperature gradually until -40°C to freeze the liquid in the PCR tubes strip for 2 hrs. Next, the chamber pressure was decreased (from 760 mTorr to 100 mTorr) to establish the primary drying phase, enabling the sublimation of all ice and the formation of a porous network. All freeze-drying phases (freezing, primary drying, and secondary drying) were programmed sequentially at fixed time points, and within each phase, critical process parameters were typically kept constant or linearly interpolated between two setpoints. The procedure was as follows: -40°C for 720 min, -20°C for 60 min, 0°C for 60 min, 10°C for 60 min, and 25°C for 480 min. The pressure of the freeze dryer chamber was maintained at less than 100 mTorr throughout the freeze-drying. Once the freeze-drying was complete, we packaged the dried mix into an aluminum foil bag using a vacuum packaging machine (DZ-400, Shanghai Hongde Packaging Machinery Co. LTD, China). The entirety of the above process was performed in an environment with a humidity of less than 3%.

### 2.4 Karl-Fischer titration

Residual moisture determination was performed on a Karl-Fischer titrator (ZDJ-2S, Beijing Xianqu Weifeng Technology Development Co., China) according to the manufacturer’s protocol. First, we cleaned the pipeline of the Karl-Fischer titrator using Karl-Fischer reagent (Sangon Biotech, China), then added the reaction buffer [50% methyl alcohol (China National Medicines Corporation Ltd.) and 50% formamide (Sigma-Aldrich)] to the reaction cup. We then weighed the freeze-dried reagents using an analytical balance (BS 224 S, 0.1 mg, Sartorius) and measured their moisture content using a calibrated Karl-Fischer titrator.

### 2.5 Sensitivity, stability, and specificity of the tests

The sensitivity of the freeze-dried PCR reagents (relative to freshly-prepared wet reagents) was tested using a 10-fold serial dilution of nucleic acid. Each reagent was reconstituted in 35 µl of nuclease-free water before adding 5 µl of the sample. We also tested how the freeze-dried PCR reagents performed if reconstituted directly in 40 µl of the sample solution. To verify the stability of the freeze-dried PCR reagents, 12 batches of SARS-CoV-2 PCR reagents were tested using a 10-fold serial dilution of nucleic acid. To evaluate the specificity, we used throat swab samples collected from five COVID-19 patients and 21 healthy controls.

### 2.6 Long-term stable test and accelerated stable test

The freeze-dried PCR mixes were stored at ambient temperature, 37°C, and 56°C, and then reconstituted to their original volume with nuclease-free water at a periodic interval. Retention of the reaction activity of the freeze-dried PCR mixes was tested (relative to freshly-prepared wet reagents) by rRT-PCR.

## 3 Result

### 3.1 Do the supplemental ingredients affect PCR performance?

To test whether the lyophilization additives had an effect on the PCR, we added trehalose [10% final concentration (w/v), Sigma-Aldrich], mannitol [1.25% final concentration (w/v), Sigma-Aldrich], BSA [0.002% final concentration (w/v), TAKARA] and PEG20000 [0.075% final concentration (w/v), Sigma-Aldrich] to the PCR mix. The amplification efficiency and cycle threshold (Ct) value were mostly unaffected by the addition of the lyophilization additives, but the fluorescence intensity (Rn) was marginally decreased (Fig. 1A–C). This indicates that the lyophilization additive had no obvious effect on PCR and could be used for subsequent lyophilization.

**Figure 1.**
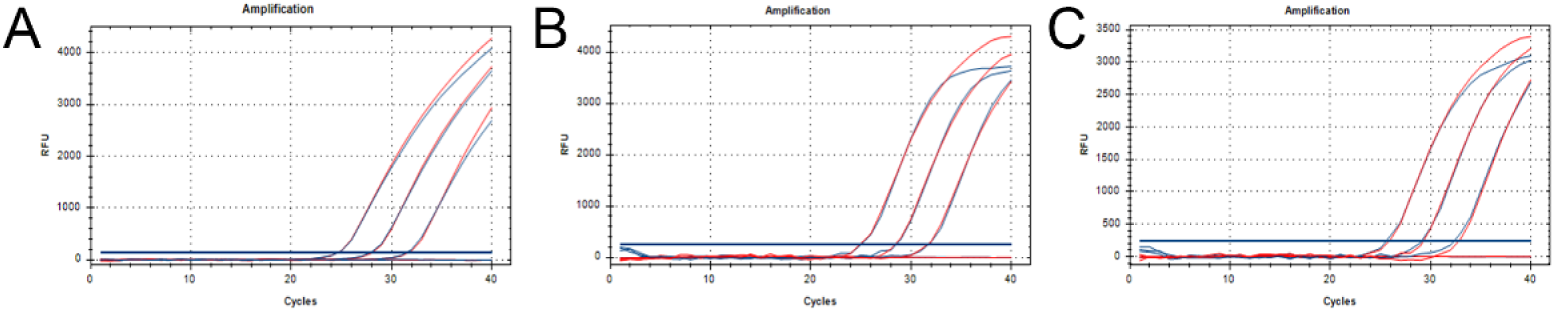
How the lyophilization additives affect the PCR. (A-C) Amplification results of the ORF1ab, N, and S genes. The red amplification curves represent the post-optimized PCR with lyophilized additives while the blue amplification curves represent the post-optimized PCR without lyophilized additives.

### 3.2 Physical appearance of the freeze-dried reagents

After lyophilization, the PCR mixes became solid with good appearance, and no obvious defects or powder diffusion were detected (Fig. 2A). To test whether the freeze-dried reagents aggregate to the edge of the PCR tubes during transportation, we placed the PCR tube strips in a regularly used vehicle for 28 days to simulate their transport by road.

**Figure 2.**
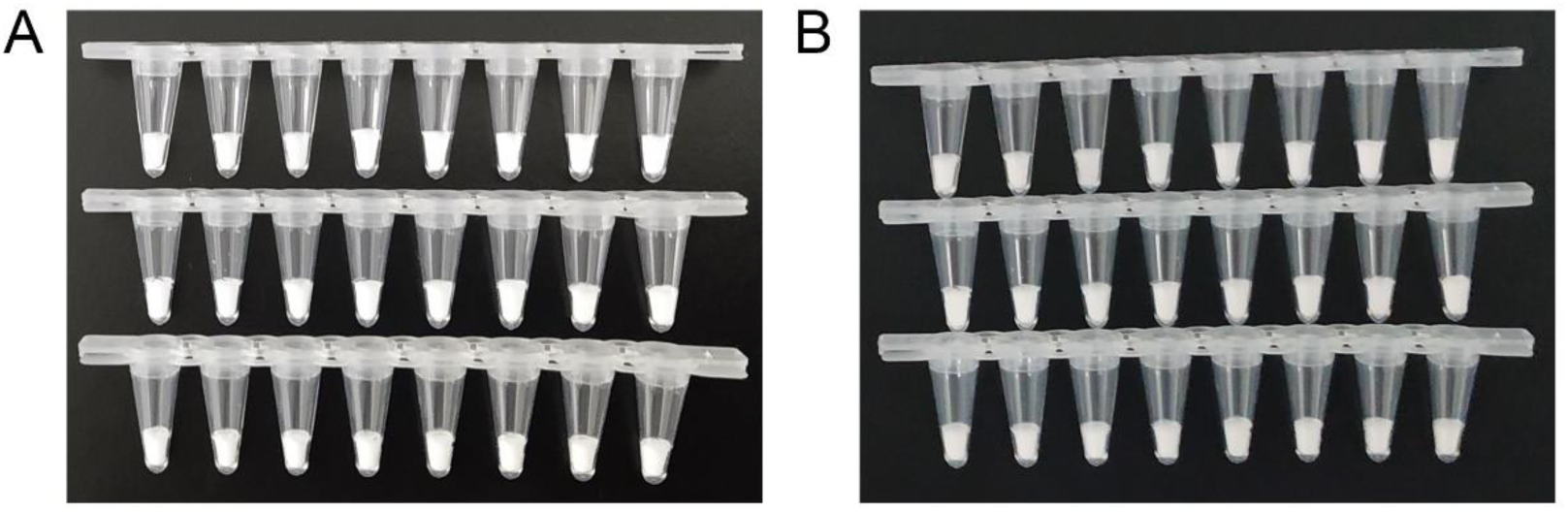
Physical appearance of the freeze-dried reagents. (A) Appearance immediately after lyophilization. (B) Appearance after simulating transportation for 28 days. From top to bottom, the freeze-dried reagents for detection of the ORF1ab, N, and S genes.

Figure 2B shows the freeze-dried PCR mixes after 28 days of simulated transport. The appearance of the reagents was unchanged by the simulated transport, and no powder floating was observed. This is likely because of the inclusion of PEG20000, a biomacromolecule that helps maintain the shape of the freeze-dried product.

### 3.3 Residual moisture content of the lyophilized reagents

Residual moisture content determination was performed on a Karl-Fischer titrator. Each set of the lyophilized mixes was measured three times with residual moisture around 1.2% (Table 2). In general, the level of addition agents in the PCR reagents and freeze-drying procedure should be adjusted to allow moisture levels of less than 3%; the residual moisture obtained by the lyophilization method presented here is appropriate.

**Table 2.** Residual moisture content of the freeze-dried PCR mixes, as measured by Karl-Fischer titration.

| Primers or probes | ORF1ab gene (%) | N gene (%) | S gene (%) |
| --- | --- | --- | --- |
| Test 1 | 1.224 | 1.197 | 1.133 |
| Test 2 | 1.242 | 1.138 | 1.261 |
| Test 3 | 1.134 | 1.183 | 1.280 |
| Mean | 1.200 | 1.173 | 1.225 |

By comparing the residual moisture of the ORF1ab, N, and S gene-targeting PCR mixes, we found that the differences among these was not obvious, and are smaller than the error caused by the measurement method itself. This indicates that the primers and probes were not major factors affecting the moisture content. Based on this finding, we propose that this method can now be transferred to other PCR mixes, changing only the primers and probes.

### 3.4 Sensitivity and repeatability of the lyophilized reagents

In these rRT-PCR assays, a 10-fold dilution series of nucleic acid was used as the reaction template. Each freeze-dried reagent was reconstituted in 35 µl of nuclease-free water before adding 5 µl of the sample, whereas the wet reagent reactions were made up of 35 µl of freshly-prepared PCR mix and 5 µl of the sample. The amplification efficiencies and Ct values were similar when comparing the freeze-dried reagent and wet reagent, while the fluorescence intensity of the freeze-dried mixes was lower than that of the wet reagent (Fig. 3A–C). Our assay was sensitive at template concentrations of 10^−5^, but not at 10^−6^ (Table 3).

**Table 3.** PCR Ct values when using various probes, before and after freeze-drying.

| Samples | ORF1ab gene |  |  | N gene |  |  | S gene |  |  |
| --- | --- | --- | --- | --- | --- | --- | --- | --- | --- |
|  | lyo | all | con | lyo | all | con | lyo | all | con |
| 10<sup>-1</sup> | 24.43 | 21.31 | 24.37 | 23.81 | 21.05 | 23.66 | 24.04 | 20.96 | 23.83 |
| 10<sup>-2</sup> | 27.98 | 24.94 | 27.86 | 26.94 | 24.29 | 27.19 | 27.12 | 24.26 | 26.72 |
| 10<sup>-3</sup> | 31.39 | 28.29 | 31.20 | 30.43 | 27.51 | 30.34 | 30.57 | 27.49 | 30.42 |
| 10<sup>-4</sup> | 34.77 | 31.73 | 34.50 | 33.71 | 31.02 | 33.70 | 33.20 | 31.21 | 33.11 |
| 10<sup>-5</sup> | 38.08 | 35.35 | 38.19 | 37.06 | 33.84 | 37.20 | 36.33 | 34.21 | 36.97 |
| 10<sup>-6</sup> | N/A | 36.68 | N/A | NA | 36.32 | NA | NA | 37.50 | NA |
| NC | N/A | N/A | N/A | N/A | N/A | N/A | N/A | N/A | N/A |

Lyo: reconstitute the freeze-dried regent with 5ul samples and 35 ul nuclease-free water; All: reconstitute the freeze-dried regent with 40ul samples completely; Con: PCR reagents without lyophilized; NC: negative control; N/A: no nucleic acid.

**Figure 3.**
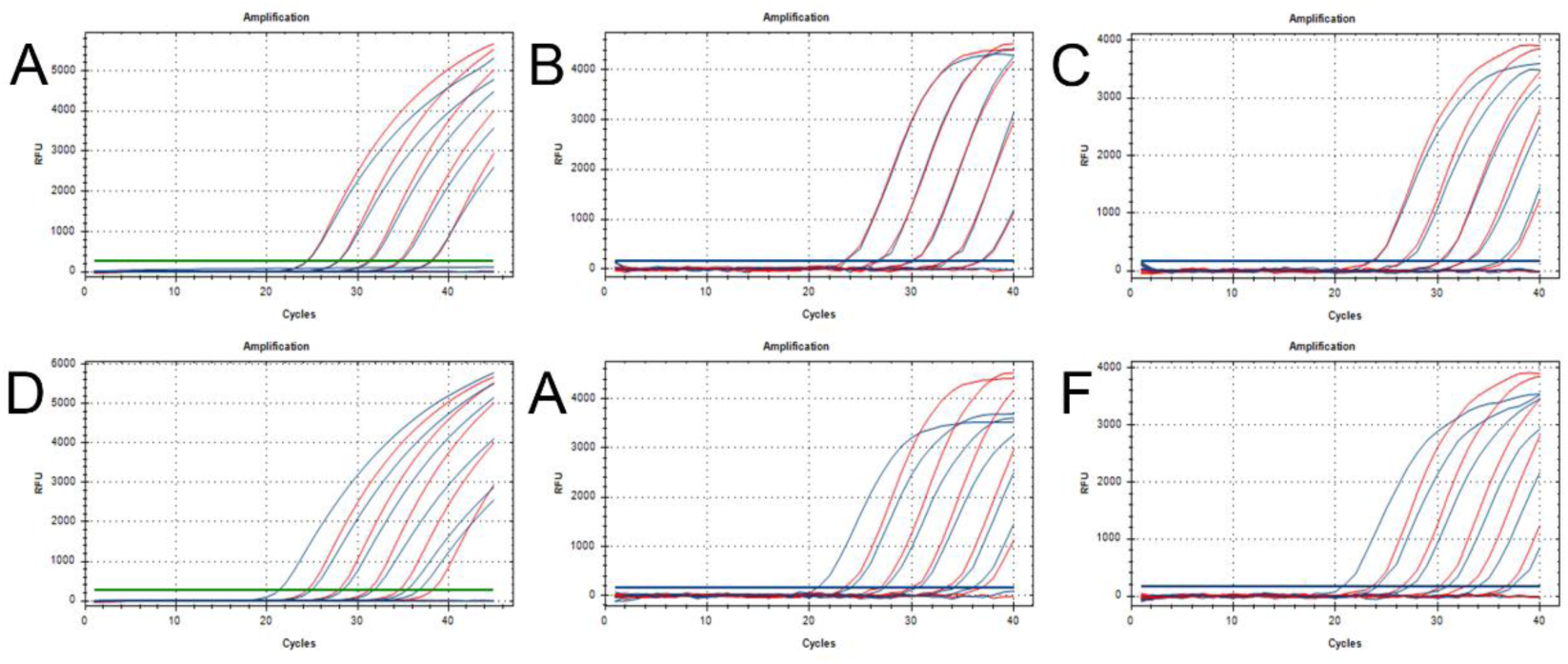
Sensitivity of the SARS-CoV-2 PCR assay using freeze-dried PCR mixes. (A–C) Amplification results for ORF1ab (A), N (B), and S (C) genes (freeze-dried vs wet reagents, the blue amplification curve represents the results with the lyophilized additives and the red line is the control without lyophilized additives). (D–F) Amplification results for ORF1ab (D), N (E), and S (F) genes (the blue amplification curves represent the freeze-dried regent reconstituted directly in 40 µl of sample solution; the red amplification curves represent the wet reagents containing 35 µl of PCR mix and 5 µl of sample solution).

To enhance sensitivity, we attempted to reconstitute the freeze-dried regent as a 40-µl total volume mix. This equates to an 8-fold increase in the sample template, which would theoretically reduce the Ct values by three. The amplification results are shown in Table 3 and Figure 3D–F. The fluorescence intensity and amplification efficiency of the former did not decrease, and the Ct values were consistent with the theoretical calculation, reduced by three. By this approach, the assay was sensitive down to template concentrations of 10^−6^.

In the repeatability assay, a 10-fold serial dilution of SARS-CoV-2 nucleic acid was selected as the reaction template, and 12 batches of lyophilized mixes were randomly selected for testing. We detected no meaningful differences in Ct value when comparing the lyophilized reagents and wet reagents (Table 4). The CV of the lyophilized reagent was larger than that of the wet reagent, but the difference was not statistically significant (P_ORF1ab_ = 0.9920; P_N_ = 0.5851; P_S_ = 0.9374, respectively). However, it is worth noting that CV tended to increase with the decrease of sample concentration in both the lyophilized group and the control group. This is determined by the characteristics of PCR detection itself, which has little relation to lyophilization. Thus, we show that the lyophilized reagents possess good repeatability.

**Table 4.** Repeatability of the PCR assay using freeze-dried reagents.

| Sam<br>ples | ORF1ab gene |  | N gene |  | S gene |  |
| --- | --- | --- | --- | --- | --- | --- |
|  | lyo(CV%) | con(CV%) | lyo(CV%) | con(CV%) | lyo(CV%) | con(CV%) |
| 10<sup>-1</sup> | 23.87(0.42) | 23.95(0.26) | 24.24(0.52) | 24.05(0.40) | 23.74(0.81) | 23.68(0.56) |
| 10<sup>-2</sup> | 27.28(0.23) | 27.38(0.18) | 27.32(0.43) | 27.24(0.39) | 26.98(1.32) | 26.83(0.78) |
| 10<sup>-3</sup> | 30.29(0.32) | 30.16(0.26) | 30.82(0.65) | 30.63(0.63) | 29.83(0.45) | 29.87(0.77) |
| 10<sup>-4</sup> | 33.90(0.94) | 33.77(0.85) | 34.30(0.59) | 34.18(0.94) | 33.45(0.81) | 33.13(1.71) |
| 10<sup>-5</sup> | 37.03(1.47) | 37.02(1.81) | 37.12(1.41) | 37.55(2.44) | 36.61(2.01) | 36.60(1.73) |
| NC | N/A | N/A | N/A | N/A | N/A | N/A |

Data are means and CV (%) for 12 groups of freeze-dried and wet PCR reagents
Lyo: lyophilization; NC: negative control; N/A: no nucleic acid.

### 3.5 Stability of the lyophilized reagents

The freeze-dried PCR mixes were stored for up to 28 days at either room temperature, 37°C, or 56°C, and, upon reconstitution, were tested relative to freshly-prepared wet reagents. At day zero, the Ct values and fluorescence intensities obtained using the lyophilization reagent were not decreased relative to the wet reagent (Fig. 4), indicating that PCR mixes could retain activity following lyophilization.

**Figure 4.**
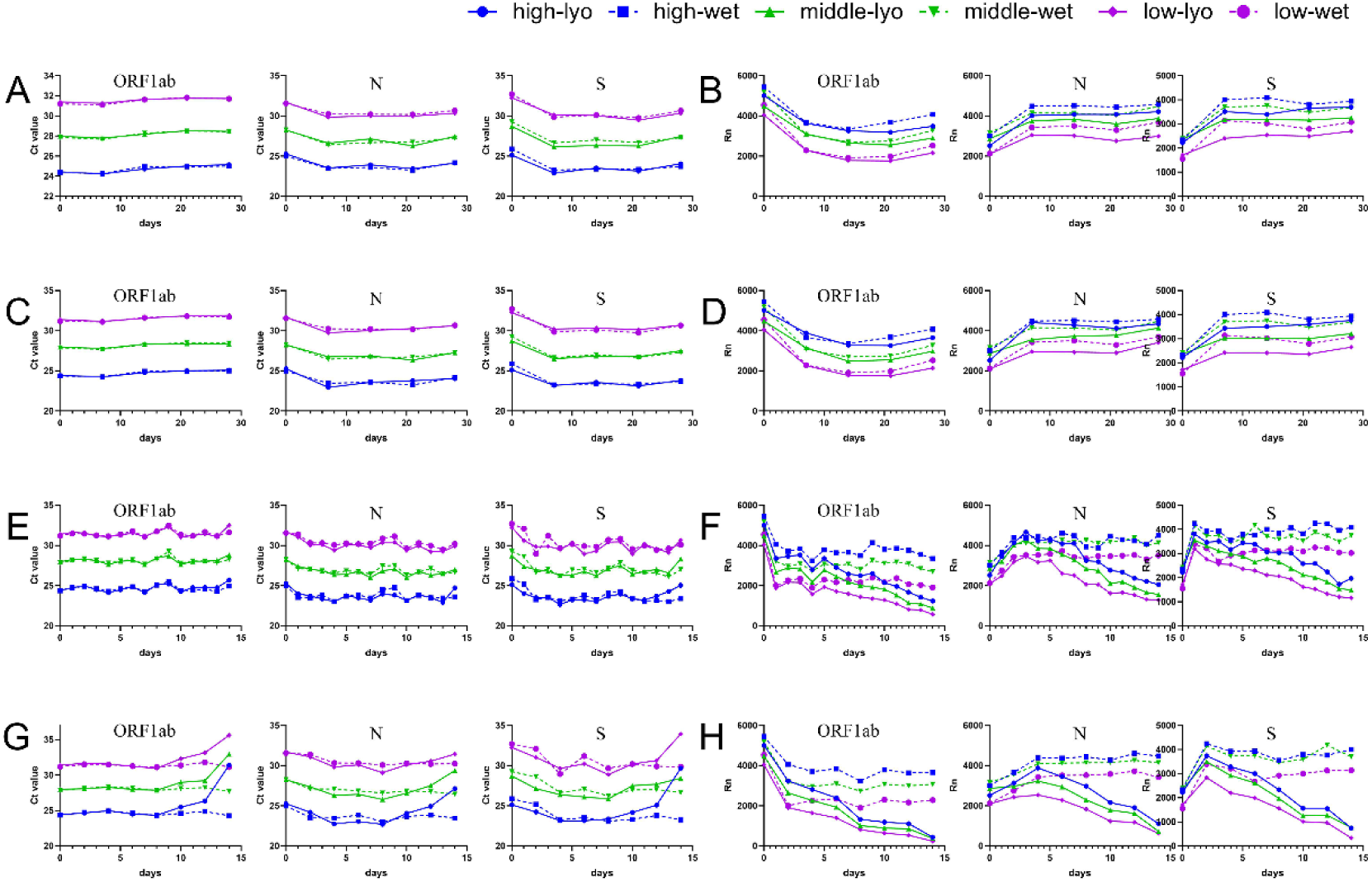
Long-term stable test and accelerated stable test of freeze-dried PCR mixes. The small pictures from left to right represent the ORF1ab, N, and S gene assays. (A) The changes in Ct values of the freeze-dried PCR mixes stored at room temperature. (B) The changes in fluorescence intensity of the freeze-dried PCR mixes stored at room temperature. (C) The changes of Ct values of the freeze-dried PCR mixes loaded on a vehicle to simulate long-distance room temperature transport. (D) The changes in fluorescence intensity of the freeze-dried PCR mixes loaded on a vehicle to simulate long-distance room temperature transport. (E) The changes in Ct values of the freeze-dried PCR mixes stored at 37°C. (F) The changes in fluorescence intensity of the freeze-dried PCR mixes stored at 37°C. (G) The changes in Ct values of the freeze-dried PCR mixes stored at 56°C. (H) The changes in fluorescence intensity of the freeze-dried PCR mixes stored at 56°C.

The freeze-dried PCR mixes were then tested at multiple time points during storage. After storing at room temperature for 28 days, similar Ct values (Fig. 4A) and fluorescence intensity (Fig. 4B) were observed for freeze-dried and wet reagents. It should be noted that the fluorescence intensity reported by the instrument fluctuates. Therefore, we use the fluorescence intensity change relative to the wet reagents as our main evaluation criterion.

We also simulated transport of the freeze-dried reagents at room temperature. After 28 days of simulated transport, the appearance of the freeze-dried mixes was unchanged (Fig. 2B). Similar Ct values (Fig. 4C) and fluorescence intensity (Fig. 4D) were observed for freeze-dried and wet reagents when targeting the ORF1ab, N, and S genes.

Ideally, we would have liked to test the activity of the freeze-dried master mix after 12 months of storage at ambient temperature. However, given the ongoing outbreak and our eagerness to share our findings, we opted to perform accelerated stability tests at 37°C and 56°C. After storing at 37°C for 2 weeks, the freeze-dried reagents performed similar (Ct values) to the wet reagents (Fig. 4E). The fluorescence intensities were initially similar, but decreased gradually from the sixth day. The mixes retained half of their original fluorescence intensity until the 14th day (Fig. 4F). When stored at 56°C, the freeze-dried reagents and freshly-prepared wet reagents initially perform similarly (Ct values), but the freeze-dried mixes lose activity from the tenth day (Fig. 4G). The fluorescence intensity values decreased sharply at the beginning, and little fluorescence could be detected on the 14th day (Fig. 4H).

In conclusion, the freeze-dried mixes retain activity at room temperature for 28 days, and for 14 and 10 days at 37°C and 56°C respectively. Also, there were no obvious differences in the results obtained for the ORF1ab, N, and S genes. This indicates that probes and primers are not the shelf-life limiting components, and that this method could be transferred to the detection of other pathogens by simply changing the probes and primers.

### 3.6 Clinical sample results

Five samples of clinical pharyngeal swabs from patients with a positive diagnosis of COVID-19 and 21 samples from healthy controls were tested using both the freeze-dried mix and freshly-prepared wet reagents. All reactions using the five patient samples tested positive (Table 6). All 21 healthy subject samples tested negative in all reactions. This indicates that the freeze-dried reagents have good specificity and can distinguish between healthy and SARS-CoV-2-infected samples, matching the performance of the freshly-prepared wet reagents.

## 4. Discussion

Freeze-drying is widely applied for the preservation and transportation of heat-labile biological drug substances at ambient temperature^[21, 22]^. In this study, we present an optimized freeze-drying formulation and procedure, allowing the stabilization of the PCR mixes at ambient temperature. We used both physical and biological methods to evaluate them comprehensively and systematically.

An ideal appearance is the basic requirement for lyophilized reagents’ packing, transportation, and preservation. It is mainly influenced by additive formulation and freezing process. Trehalose, as an important lyophilization protectant, and plays a crucial role in the lyophilization process. However, if the trehalose concentration is too high, the appearance of the final product can be compromised. If some macromolecular substances (e.g., PEG20000) are added in the PCR mixes, the mixed reagents can become tightly connected after lyophilization, which can help to avoid disturbance during transportation. The freezing process of freeze-drying can be divided into three stages, and it is important to ensure that the reagents can be maintained at low temperatures for sufficient time during the freezing process to make sure that ice crystals can grow to the extent that no further ice crystal growth is possible. Otherwise, the appearance of the freeze-dried reagents may be affected^[23]^.

Residual moisture content is an impact factor influencing the quality and stability of freeze-dried PCR mixes^[18, 19]^. A high moisture level will decrease the stability of the reagent. Since glycerol is hygroscopic, its presence in the final freeze-dried product likely results in a high moisture content, which might affect the stability of the product^[24]^. The commercial availability of glycerol-free Taq polymerases (enzyme) would help to prolong the shelf life of freeze-dried PCR mixes^[11]^. However, removing all water from the reagent would have deleterious effects on those reaction components, proteins for example, that require certain amounts of bound water in order to maintain proper conformations. Here, we found that a residual moisture content of 1–3% is optimal for freeze-dried PCR mixes.

We chose rRT-PCR to evaluate the detection performance of the freeze-dried PCR mixes. The supplemental ingredients added to the freeze-dried mixes did not affect the Ct values, fluorescence intensity, or amplification efficiency of the PCR mixes. The sensitivity, specificity, and repeatability of freeze-dried reagents were similar to those of the freshly-prepared wet reagents. We also found that the sensitivity of freeze-dried PCR mixes could be improved by reconstituting the dried mix using the test sample solution (to a volume of 40 µl). However, we did not observe the activity of lyophilized PCR mixes beyond 28 days of storage. Given the ongoing outbreak and our eagerness to share our findings, we opted to use an accelerated stability test to predict the long-term storage effect of the lyophilized reagent at room temperature. Klatser et al. described a freeze-dried PCR mix for detection of mycobacteria, which could retain activity at 4°C and 20°C for 1 year and at 56°C for 1 week^[11]^. Unlike in the work of Klatser et al., our freeze-dried PCR mixes contain a reverse transcriptase. Given that our freeze-dried PCR mix could retain activity at 56°C for 10 days, we predict that it would remain active for 1 year when stored at room temperature.

In conclusion, we describe a method for producing thermostable freeze-dried PCR mixes for use in COVID-19 diagnosis, with sensitivity, specificity, and repeatability values that match those of freshly-prepared wet reagents. There were no obvious differences in the performance of the freeze-dried mixes targeting the ORF1ab, N, and S genes of SARS-CoV-2. Based on this finding, we propose that the primers and probes do not affect the efficiency of the lyophilization.

We propose that the method described here can now be transferred to the lyophilization of PCR mixes targeting other pathogens by simply changing the primers and probes. This approach will also be useful in tackling future major outbreaks or other public health hazards.

## Acknowledgements

This work was supported by the Xiamen Science and Technology Major Project (Grant No. 3502Z2020YJ01), the National Science and Technology Major Project of China (Grant No. 2018ZX10732101-001-002), and the National key research and development program (Grant No. 2018YFC1200103).

